# An AI for an AI: identifying zoonotic potential of avian influenza viruses via genomic machine learning

**DOI:** 10.1101/2025.09.16.676011

**Authors:** Liam Brierley, Joaquin Mould-Quevedo, Matthew Baylis

## Abstract

Avian influenza remains a serious risk to human health via zoonotic transmission, as well as a feasible pandemic threat. Although limited zoonotic cases have resulted from the current epizootic outbreak, the wide diversity of influenza viruses in avian hosts means the emergence of new strains that could transmit to humans more readily cannot be ruled out. There is therefore a need to anticipate zoonotic potential before spillover occurs.

Here, we develop a novel zoonotic prediction model for avian influenza viruses, building upon “host-predictor” machine learning methods that estimate host potential given only a viral genome sequence. We construct a machine learning framework combining individual sub-models of influenza genome segments, each trained on many genomic and proteomic traits (e.g., k-mer composition, codon biases, protein physicochemistry).

To prevent over-fitting to heavily sampled lineages and ensure models generalise to phylogenetically distant viruses, we pre-process training data by considering clusters of shared sequence identity. Curated training sets cover ∼4,000 representative, complete genome sequences of avian influenza from 120 subtypes including 9 containing known zoonotic viruses.

We combine best-performing models into a single ensemble that can distinguish zoonotic capability of sequences held out from training with strong performance (AUROC = 0.95, F1 score = 0.90), including sequences of rarely-sampled subtypes, e.g., H10N8. Interrogating ensemble model decisions also allows us to identify influential genomic motifs most associated with human infection.

These findings suggest specific genomic traits that are key to understanding and monitoring evolution of influenza viruses that circulate within bird populations. Our ensemble model can estimate zoonotic potential for new sequence inputs, offering a means to quickly risk-assess emerging avian influenza strains as soon as a sequence becomes available.

## Introduction

Avian influenza is a critically important wildlife-origin disease for impacts on poultry industries as well as for potential human pandemic risks. Diversity among avian influenza A viruses is maintained by populations of wild birds, primarily waterfowl, and many different subtypes can co-circulate globally. Since 2020, the 2.3.4.4b clade of H5 highly pathogenic influenza has sparked a wide epizootic in wild birds (Xie et al. 2023) with many resulting incursions into mammals, a stark reminder that influenza viruses are frequently capable of crossing species barriers. Infection has been identified in species including cats (Domańska-Blicharz et al. 2023), mink (Agüero et al. 2023), and wild carnivores such as foxes, otters, and lynx (Tammiranta et al. 2023). These have mostly been restricted to dead-end infections, though two notably large mammalian outbreaks reignited concerns around mammal-to-mammal transmissibility. Firstly, mass mortality events were seen in populations of marine mammals (e.g., sea lions, elephant seals) in South America (Puryear et al. 2023, Leguia et al. 2023, Ulloa et al. 2023), with evidence of mammalian-like adaptation in the PB2 gene (Rimondi et al. 2024). Infection has also been increasingly found in dairy cattle across North America (Burrough et al. 2024, Ly 2024) with potential mammal-to-mammal transmission via contact with raw milk in shared milking equipment (Sage et al. 2024). Subsequent occupational infection of farm workers has prompted high-priority surveillance responses (Garg et al. 2024).

In total, zoonotic spillover of clade 2.3.4.4b H5N1 has so far resulted in a modest number of identified human cases. Most were occupational infections with minimal clinical presentation (Webby and Uyeki 2024), though at least one fatality has been recognised. However, zoonotic transmission of avian influenza has been a recurrent public health challenge far beyond clade 2.3.4.4b. Zoonotic infection has been detected for avian influenza viruses spanning fifteen subtypes to date (see Liu et al. (2022) for an earlier inventory), most recently H3N8 in Hunan, China in 2022 (Yang et al. 2022) and H5N2 in Mexico City, Mexico in 2024 (Vázquez-Pérez et al. 2024). Reported case fatality ratios have varied from 39 – 61% across these incidents (World Health Organisation 2024), though these figures are likely over-estimates from biases against detecting mild or asymptomatic cases. No strong evidence for human-to-human transmissibility has yet been observed, although emergence of a more transmissible zoonotic avian influenza virus is considered a feasible and urgent pandemic threat (Burke and Trock 2018, Yamaji et al. 2024), especially considering the potential for adaptation during circulation in mammalian livestock.

Anticipating the likelihood of humans (or any other species) being a novel host for a given virus is now more feasible following developments in computational genomics. Various methods can estimate host suitability by leveraging the complex, multidimensional information within virus genome sequences (Mollentze and Streicker 2023). In practice, this requires careful training of machine learning algorithms to recognise implicit patterns in compositional, structural, or physiochemical features of - omics data that correlate with host range phenotypes. Resulting patterns can be generalised to make predictions even on highly divergent ‘out-of-sample’ sequences. Much of the work in this accelerating field has used sequences to either estimate high-level host taxa, e.g., prediction of source host orders, classes, or families (Babayan et al. 2018, Young et al. 2020, Brierley and Fowler 2021) (but see Mock et al. (2021) for a counter-example), or estimate whether a high-level breadth of viruses will infect a single given host species (Young et al. 2020, Bartoszewicz et al. 2021, Mollentze et al. 2021). Influenza A viruses feature a negative-sense single-stranded RNA genome composed of eight separate protein-encoding segments, which allows for segment reassortment. We assert that influenza host predictor models need to accommodate for this segmentation and here we focus on models tailored to the broad diversity of influenza A viruses.

Most existing host predictor models of influenza A have discriminated between viruses that are relatively established within avian, swine, and/or human populations (e.g., seasonal human influenza) (Yin et al. 2019, Li et al. 2020, Mock et al. 2021, Xu and Wojtczak 2022) using a variety of sequence-derived features (Borkenhagen et al. 2021). Anticipating zoonotic transmission, however, requires more bespoke analyses as the genomic signals associated with capability for spillover infection of a human host are likely to be distinct from those that result from longer-term host adaptation (Greenbaum et al. 2008, Long et al. 2019).

Several previous studies have specifically trained models to predict zoonotic infection from avian influenza sequences. Eng et al. (2017) trained random forest algorithms on sequences covering a broad diversity of influenza A subtypes, including six subtypes containing zoonotic viruses. Their models featured composition of protein motifs based on amino acid properties, which can act as a proxy for protein structure and subsequently, for host-virus interactions. They trained models on each protein, before combining into one summative model to distinguish sequences with zoonotic potential (Eng et al. 2017). Sun et al. (2021) carried out a mutational analysis of the H7N9 subtype by considering nucleotide identity at each position in whole genome alignments as model features. Their logistic regression is one of the few prediction models to be validated in a lab setting, finding selected viruses predicted to be zoonotic were more infective in mammalian cell line and live mouse inoculations (Sun et al. 2021). Similarly, Scarafoni et al. (2019) predicted zoonotic capability from amino acid identity at each position in alignments of several proteins across multiple influenza A subtypes. They designed a convolutional neural network (cNN), a sophisticated approach that can determine its own features through densely connected filtering layers. Each study was then able to suggest likely strains or lineages for future zoonotic spillover by applying models to novel data.

However, none of these approaches explicitly adjusted the training methodology for sequence relatedness. Model performance can be inflated by training and testing on phylogenetically related genome sequences, as model-learned patterns may simply reflect membership of the same lineage or subtype rather than true molecular host compatibility (Di Giallonardo et al. 2017, Mollentze and Streicker 2023). Scarafoni et al. (2019) found that when retrained on subtype membership, model performance was almost as strong as that of the sequence-informed cNN, demonstrating high potential for this bias in influenza A datasets. This overfitting to phylogenetic structure can then lead to failure to generalise to novel sequences, particularly if they are phylogenetically distant.

Here, we build on previous works to develop a zoonotic prediction model tailored to the influenza A genome that incorporates a much wider range of features capturing both nucleotide and protein traits that do not require prior alignment. We use a segment-by-segment training strategy combined into an overall ensemble to distinguish zoonotic avian influenza (from human samples, covering nine subtypes) from non-zoonotic avian influenza (from wild or domestic birds). Critically, we attempt to mitigate phylogenetic bias by pre-processing our training data based on sequence clustering to ensure predictions are generalisable. By developing a system to give reliable zoonotic predictions on unseen data, our models represent an inexpensive and rapid risk assessment process for newly emerging avian influenza lineages.

## Methods

### Data extraction

We identified and extracted complete whole genome sequences of influenza A virus (defined as those featuring complete sequences for all 8 segments) from the NCBI Influenza Virus Resource (Bao et al. 2008) and the GISAID repository (Shu and McCauley 2017), taking all available sequences at date of extraction (21/02/23). We used metadata host labels to assign sequences as zoonotic if they were sampled from humans and had a H5, H6, H7, H9, H10 subtype or a H3N8 subtype. We assigned sequences as avian (as a proxy for non-zoonotic) if they were sampled from any wild or domestic bird species, regardless of subtype. Duplicate sequences and those from mixed-subtype infections were discarded. These filters resulted in 618 zoonotic whole genomes and 18,913 avian whole genomes, collectively covering 120 subtypes (Fig. S1).

To remove redundancy and thin representation of heavily-sampled lineages, we then conducted sequence clustering using Linclust (linear-time clustering) within the MMseqs2 software. Computation time of this algorithm scales linearly with size of input, benefiting clustering tasks on large sequence sets (Steinegger and Söding 2018). Linclust groups sequences based on shared k-mers, before locally aligning to link specific sequences within-group. We considered sequences linked if sharing at least 70% identity (min-seq-id = 0.7) over 70% mutual coverage in alignment (c = 0.7). Link structure is then used to determine final clusters via a greedy incremental algorithm (Hauser et al. 2016). Initial clustering of complete sequence data generated 2018 clusters, for each of which we retained only the most central sequence to represent the cluster in model training.

This approach also allowed us to mitigate a common source of bias in analyses predicting host labels derived from metadata (Borkenhagen et al. 2021). Any influenza sequence sampled from an avian host cannot strictly be considered a ground-truth zoonotic negative, as they may have unobserved zoonotic potential or may even have originated a known zoonotic event (which cannot be determined without in-depth epidemiological study). Therefore, we did not select any avian sequences as a cluster representative if the cluster contained a mix of avian and zoonotic sequences (n = 68 clusters, 3.4%), selecting a random zoonotic sequence instead.

### Feature calculation

We generated multiple sets of features describing genomic and proteomic properties of sequences based on interpretability and suitability in previous studies (Eng et al. 2014, 2016, Yin et al. 2019, Young et al. 2020, Borkenhagen et al. 2021). Features were calculated independently for each individual segment nucleotide sequence, coding sequence (cds), and protein sequence. For the latter, we focused only on the longest ORF of each segment and its corresponding protein (e.g., M1 protein for segment 7, NS1 protein for segment 8).

For nucleotide sequences of entire segments, we calculated simple frequency of overlapping k-mers with 2 ≤ k ≤ 6. For coding sequences, we calculated multiple measures of genome composition bias following Babayan et al. (2018) and Brierley and Fowler (2021): simple nucleotide proportion; dinucleotide bias (calculated separately at each codon position, i.e. 1-2, 2-3, or 3-1, and adjusted for expectation based on nucleotide composition), codon bias (Relative Synonymous Codon Usage (Sharp and Li 1987)), and simple amino acid proportion. Compositional features were calculated using R packages ‘Biostrings’, v2.74.1 (Pagès et al. 2024) and ‘coRdon’ v1.24.0 (Elek et al. 2024).

For protein sequences, we calculated a simple initial feature set of proportions of overlapping 2-mers (dipeptide composition, or DPC). Following this, we calculated several further feature sets capturing functional protein physicochemistry. Firstly, we used Conjoint Triad features (CTriad) (Shen et al. 2007). These divide amino acids into seven groups or ‘classes’ based on overall physicochemical similarity (primarily volume and electrostatic interactions), effectively creating a reduced alphabet. Standardised frequency of each 3-mer with this alphabet is then calculated to give 7^3 features.

Secondly, we used Composition-Transition-Distribution (CTD) features (Dubchak et al. 1995, 1999). These similarly use a reduced alphabet, dividing amino acids into three groups based on physicochemical properties, though separate features are calculated for each individual property. For example, for electrostatic charge, three Composition features were calculated as simple proportion of each group (either positively charged, negatively charged, or neutral). Three Transition features were calculated as simple proportion of each 2-mer featuring adjacent non-identical groups (positive-negative, positive-neutral, negative-neutral), independent of sequence order. Lastly, five Distribution features captured proportion of sequence covered before reaching a given quartile of amino acid content from a given group, e.g., if a quarter of all positive residues present are within the first 8% of the sequence, then D_0.25_positive = 0.08. Distribution features were calculated for each group, and for the first, second (median), and third quartiles as well as the first and last residues. CTD features were calculated based on 13 groupings capturing 7 different properties.

The final feature set generated used Pseudo-Amino Acid Composition (PseAAC) (Chou 2001), which gives standardised frequences of the 20 amino acids but with λ additional features capturing cross-residue correlation. For each pair of amino acids n residues apart, mean absolute distance between physicochemical property values (e.g., mass, hydrophobicity) is calculated, before combining as a weighted average across all pairs (Chou 2001). Independent features can be then generated for values of n from 1 through λ, where we consider λ = 3 (i.e., correlation between amino acids three residues apart at most).

While CTriad features represent only composition of amino acids, CTD and PseAAC features also represent relative position of amino acids within the sequence, which are likely to better capture local structural effects of the resulting protein. All protein features were calculated using Python package ‘iFeatureOmega’ (Chen et al. 2022).

### Model training

We used each of the 12 feature sets calculated on each respective segment, cds, and protein to classify sequences as either avian or zoonotic (Fig. 1) by training five different supervised learning algorithms on each combination. We selected algorithms informed by performance in previous influenza genome classification tasks (Borkenhagen et al. 2021): penalised logistic regression (PLR) using package ‘glmnet’ v4.1-6 (Tay et al. 2023); random forests each comprising 1000 trees, using package ‘ranger’ v0.14.1 (Wright and Ziegler 2017), linear and radial support vector machines (SVM) using packages ‘e1071’ v1.7-16 (Meyer et al. 2024) and ‘kernlab’ v0.9-33 (Karatzoglou et al. 2024); and XGBoost over 2000 boosting iterations using package ‘xgboost’ v1.7.6.1 (Chen et al. 2024). Before training, all features were centred and rescaled to 0-1 and we adjusted for class imbalance by weighting data inversely proportional to class frequency, effectively upweighting training towards classifying zoonotic sequences. Model parameters were optimised by five-fold cross-validation, selecting parameters that maximised Cohen’s Kappa metric. All models were trained in R v4.2.2 using ‘caret’ v6.0-94 (Kuhn 2022).

**Figure 1.**
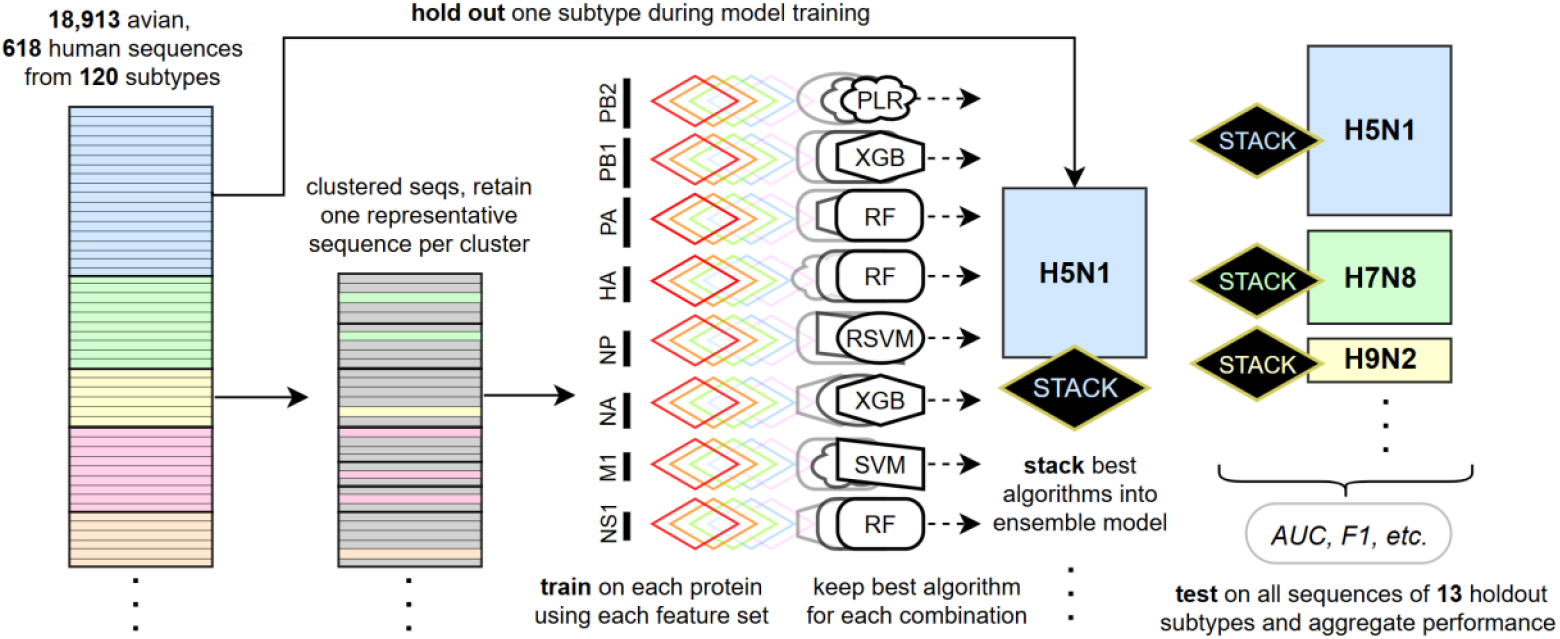
Structured machine learning model training over feature sets and influenza subtypes. Diagram indicating data partitioning and machine learning model training procedure. Individual sequences are indicated by horizontal bars, while colour indicates influenza HN subtype with greyed-out sequences being excluded before training. Coloured diamonds represent 12 calculated feature sets and shapes represent application and selection of best classification algorithm for each feature set-gene or feature set-protein combination (PLR = penalised logistic regression, RF = random forest, SVM = linear support vector machine, RSVM = radial support vector machine, XGB = XGBoost; each optimised by five-fold cross-validation). Diamonds indicate ensemble stack models, each constructed holding out a different influenza HN subtype.

Initial training of each classification algorithm upon each feature set-segment combination produced 12*8*5 = 480 independent models, which we condensed into a single ensemble interface to generate overall predictions of zoonotic risk. Firstly, we kept only the best-performing algorithm from the five in each case (Fig. 1). We then combined the resulting 96 models by training a ‘stack’, or meta-learner. This method takes the output probabilities from a set of existing models and uses them as input features to train a new model. We train a LASSO logistic regression as the stack algorithm, which penalises coefficients toward zero, effectively conducting model selection by minimising the influence of poorly-performing or redundant models. Stack models were trained using R packages ‘glmnet’ v4.1-6 (Tay et al. 2023) and ‘caretEnsemble’ v2.0.3 (Deane-Mayer 2023), optimising the lambda regularisation parameter by five instances of five-fold cross-validation.

To validate whether models could generalise to predict zoonotic status for unseen, divergent sequences, we selected 13 influenza subtypes a priori as representative test sets: 9 zoonotic subtypes with at least one whole genome sequence sampled from humans (H3N8, H5N1, H5N6, H7N3, H7N4, H7N7, H9N2, H10N8), and 4 non-zoonotic avian subtypes that did not share a haemagglutinin subtype with any zoonotic events, each having at least 150 whole genome sequences (H4N6, H4N8, H8N4, H16N3). We held out each in turn (Fig. 1) and re-clustered training data in each case (giving between 1746 and 2123 clusters depending on subtype held out) before re-training all individual models and re-training a new stack ensemble that was then applied to generate predictions on the held-out sequences.

For non-zoonotic held-out subtypes, test predictions were generated for all sequences. However, for zoonotic subtypes, as previously outlined we have less ground truth confidence in non-zoonotic sequence labels given that we cannot distinguish which were ancestral to zoonotic transmission events. Therefore, we generated predictions for confirmed zoonotic sequences only. Predictions of all 13 holdout sets were aggregated to calculate final model performance metrics including area under receiver operating curve (AUROC) and F1 score. We selected the optimal classification threshold for probabilistic predictions as the threshold giving the closest point to top left in the ROC, i.e., minimising (1 – sensitivity)^2^ + (1 – specificity)^2^.

### Model interpretation

To identify which genomic and proteomic features were most influential in distinguishing zoonotic sequences, we used a method-agnostic permutation approach to calculate variable importance. We randomly permuted each of the individual feature columns of each segment, cds, or protein in turn across all holdout sets before inputting to stacked models and again predicting zoonotic capability. We then calculated the resulting loss in performance associated with breakdown of each feature in turn, as measured by difference in AUROC pre-and post-permutation.

## Results

We extracted whole genome sequences of avian influenza spanning a period of 1902 – 2022, though over 97% of sequences were dated post-2000 (Supplemental Fig. S1). H7N9 was the dominant subtype among the extracted zoonotic sequences, with most of these originating between 2013 and 2016 (Supplemental Fig. S1, Supplemental Table S1), followed by many H5N1 sequences from 2003 to 2012. Although 120 subtypes were represented in total, H3N8, H4N6, H5N1, H5N2, and H9N2 subtypes had the highest coverage of non-zoonotic avian sequences (Supplemental Table S1).

Following sequence clustering, we trained machine learning algorithms to classify zoonotic sequences based on individual influenza virus segments. Zoonotic status among unseen test subtypes was generally better distinguished using features capturing properties of protein sequences compared to nucleotide sequences (Fig. 2), a pattern that held for both AUROC and F1 scores (Supplemental Fig. S2, S3). Specific segments of influenza virus also offered stronger predictive power for zoonotic infection. AUROC was generally highest using haemagglutinin (HA), nucleoprotein (NP), matrix protein 1 (M1), and non-structural protein 1 (NS1) genes/proteins, regardless of feature set used. Random forests and XGBoost tended to outperform other algorithms and were most often retained as the best model method (Fig. 2), particularly for larger sets of features (e.g., nucleotide 5-mers and 6-mers and CTD-D).

**Figure 2.**
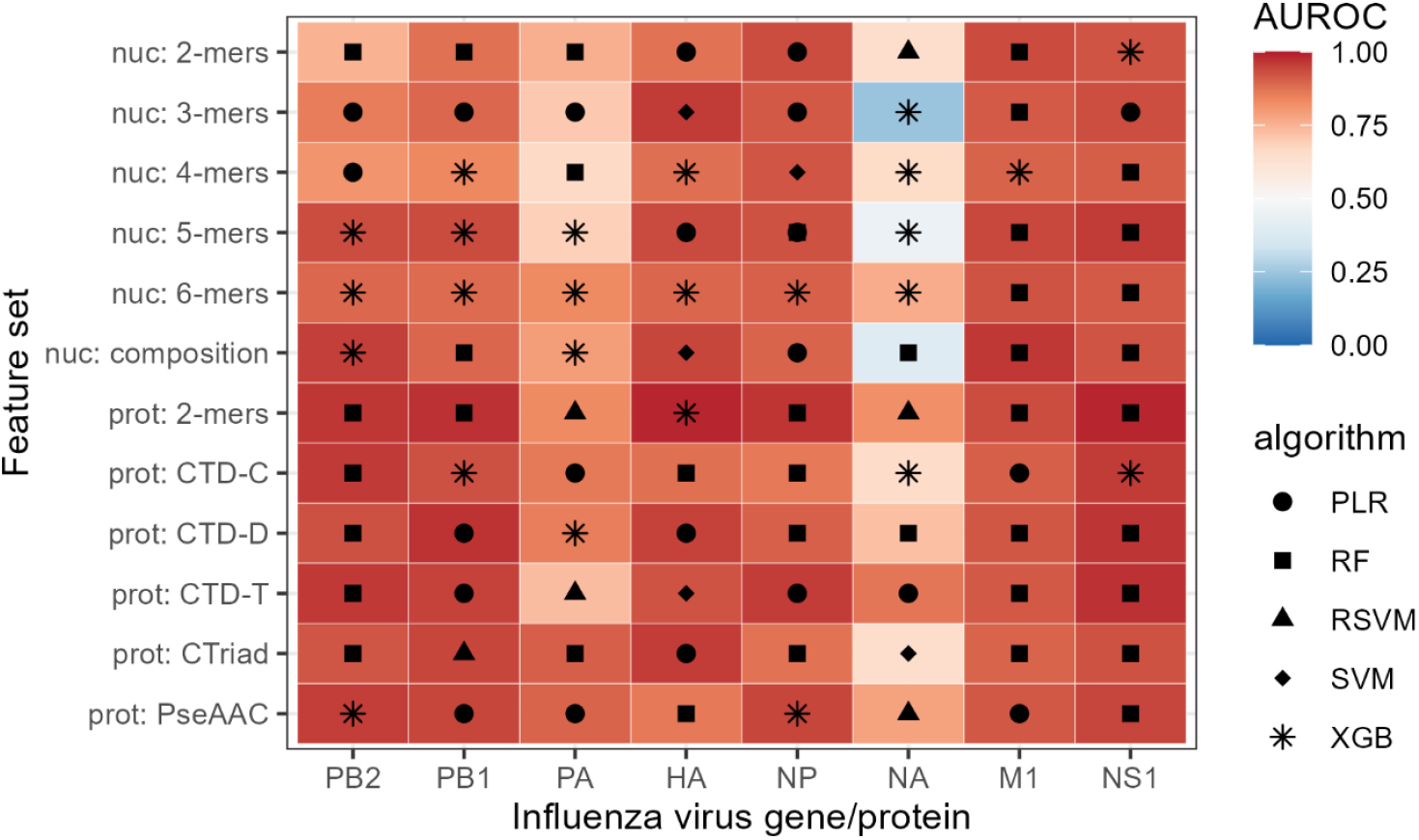
Best machine learning model performance for individual gene/proteins and feature sets. Heatmap of performance for 96 machine learning models covering each combination of influenza virus gene or protein (columns) and sequence-derived feature sets (rows). Colour scale denotes collective performance over each of the thirteen held-out test subtypes aggregated into a single measure of AUROC. Performance is only shown for the best-performing algorithm, indicated by symbol (PLR = penalised logistic regression, RF = random forest, RSVM = radial support vector machine, SVM = linear support vector machine, XGB = XGBoost). ‘nuc’ denotes nucleotide sequence features, ‘prot’ denotes protein sequence features: dipeptide composition (2-mers), Composition-Transition-Distribution (CTD), Conjoint Triad (CTriad), and Pseudo-Amino Acid Composition (PseAAC).

We then combined the 96 best-performing models (Fig. 2) into a single ‘stack’ ensemble. By collating information learned from genomic features of each individual segment, the stack method predicts zoonotic capability with strong performance (AUROC = 0.947, F1 = 0.896) and well-balanced sensitivity (0.919) and specificity (0.960). This performance generalises well to sequences from out-of-sample, unseen influenza subtypes (Fig. 3), though more variability in final stack predictions was observed for certain subtypes, e.g., zoonotic H7N9. Among five subtypes that have less frequently been reported to cause zoonotic infections, the stack method most confidently predicted available sequences of H10N8 as zoonotic, followed by a recently recognised zoonotic sequence of H3N8 from 2022 (Fig. 3). Only one rare zoonotic subtype (H7N3) was misclassified as non-zoonotic by the stack.

**Figure 3.**
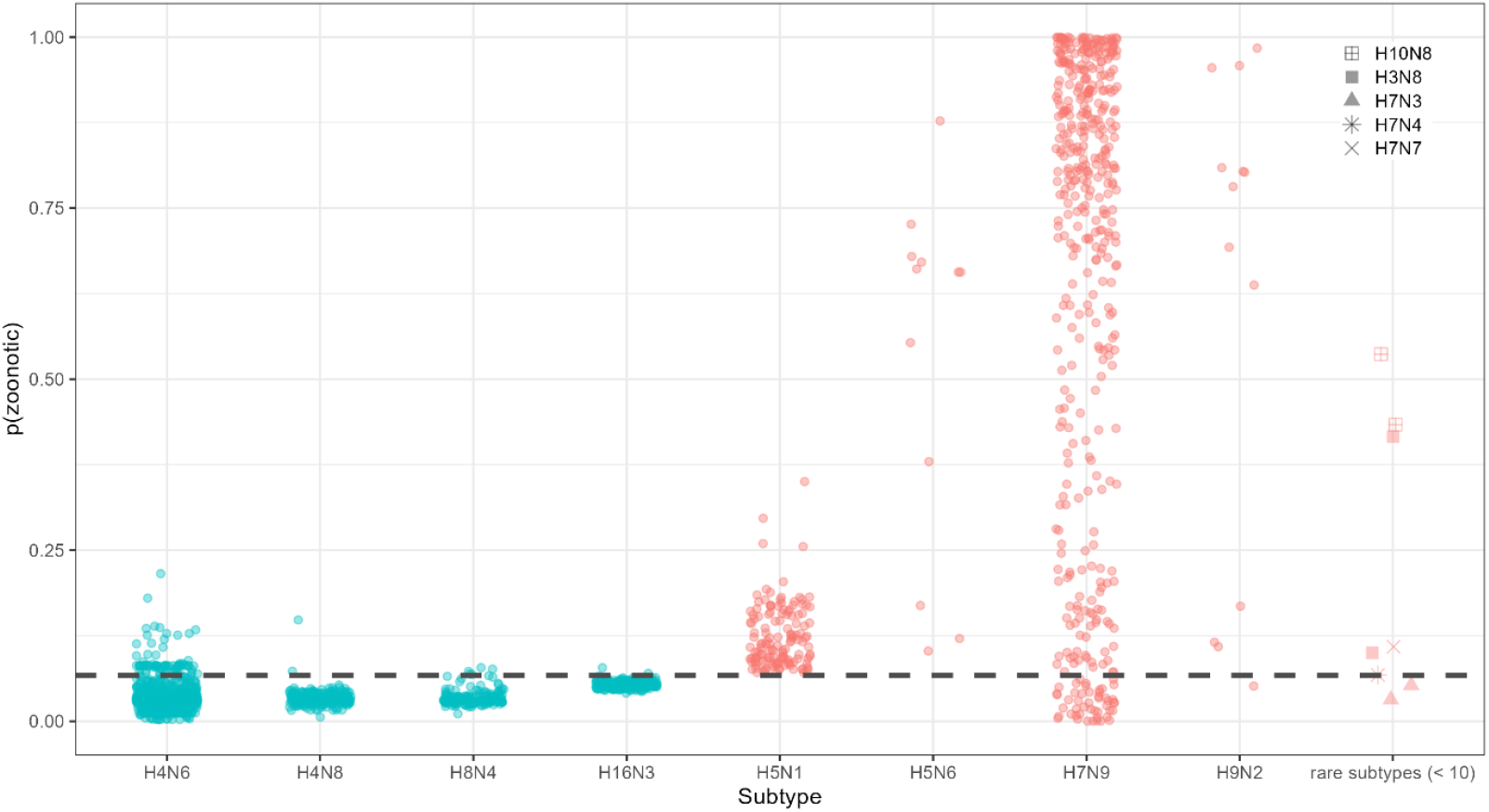
Predicted zoonotic probability for unseen test sequences held out from stack models. Stack method-predicted probability of zoonotic potential for individual sequences among the thirteen holdout subtypes. Teal blue denotes sequences from exclusively non-zoonotic avian subtypes while red denotes zoonotic sequences from known zoonotic subtypes. Rare subtypes with fewer than 10 sequences are grouped, with shape denoting exact subtype. Dashed line indicates optimal model decision cutoff (i.e., sequences with probability above line are predicted more likely zoonotic than non-zoonotic).

Generally, the stack method predicted non-zoonotic avian sequences with greater confidence, although many sequences from exclusively avian subtypes were close to or over the optimal classification threshold (Fig. 3). Based on their genomic and proteomic properties, two sequences of H4N6 and one sequence of H4N8 appeared distinctly higher in predicted zoonotic potential, all three originating from waterfowl samples in the Americas: *A/American black duck/New Brunswick/00499/2010, A/yellow-billed teal/Argentina/CIP051-91/2011*, and *A/American black duck/New Brunswick/02375/2007*, respectively.

We then investigated exactly which genes/proteins and features informed these decisions of the stack ensemble. Firstly, we examined which of the 96 individual models (Fig. 2) were retained after stack model selection. Models based on HA were most represented (Supplemental Fig. S4), with NP, NS1, and polymerase basic protein 2 (PB2) models also being well-represented.

We then interrogated which specific genomic and proteomic features had the most influence within the stack by permuting each individual feature in turn. We found ability to classify zoonotic sequences was disrupted by permuting features of each gene and protein except for M1 (Fig. 4), and only minimally so for neuraminidase (NA). This implies that zoonotic potential has wide dependency throughout the influenza genome. We also found 3-mer motifs of nucleotide sequences and physicochemical traits of protein sequences to be most informative (Fig. 4), particularly those capturing transitions between amino acid groupings, i.e., CTD-T features. These features often involved solvent accessibility or hydrophobicity (Table 1), suggesting they may be capturing elements of molecular binding processes.

**Table 1.** Top twenty avian influenza virus gene/protein features by permutation variable importance. Variable importance of twenty most influential features, measured by reduction in performance (loss in AUROC) when individual features were permuted. Influenza virus gene/protein and feature set are given, along with value of AUROC loss.

| Gene/protein | Feature set | Feature | AUROC loss |
| --- | --- | --- | --- |
| NS1 | nuc: 3-mers | TAG | 0.040 |
| NS1 | nuc: 3-mers | AGG | 0.039 |
| PB2 | nuc: 3-mers | CCT | 0.037 |
| PA | prot: CTriad | Group 3/Group 6/Group 3 triad | 0.032 |
| NS1 | nuc: 3-mers | TAA | 0.031 |
| NS1 | nuc: 3-mers | ATT | 0.030 |
| NS1 | nuc: 3-mers | CGG | 0.029 |
| PB2 | nuc: 3-mers | CCA | 0.029 |
| PB1 | prot: CTD-T | Polar-hydrophobic transition<br>(Engelman Hydrophobicity index) | 0.028 |
| NS1 | nuc: 3-mers | TTA | 0.024 |
| NS1 | nuc: 3-mers | ACA | 0.023 |
| NS1 | nuc: 3-mers | AGC | 0.021 |
| PA | prot: CTriad | Group 3/Group 4/Group 3 triad | 0.018 |
| PB1 | prot: CTD-T | Buried-exposed transition<br>(solvent accessibility) | 0.016 |
| **PB1** | prot: CTD-T | Polar-hydrophobic transition<br>(Casari-Sippl Hydrophobicity index) | 0.016 |
| **HA** | nuc: 2-mers | TG | 0.015 |
| **PB1** | prot: CTD-T | Polar-neutral transition<br>(Argos Hydrophobicity index) | 0.015 |
| **NS1** | nuc: 3-mers | TAT | 0.013 |
| **NS1** | nuc: 3-mers | TTT | 0.013 |
| **PB2** | nuc: 3-mers | GCT | 0.013 |

**Figure 4.**
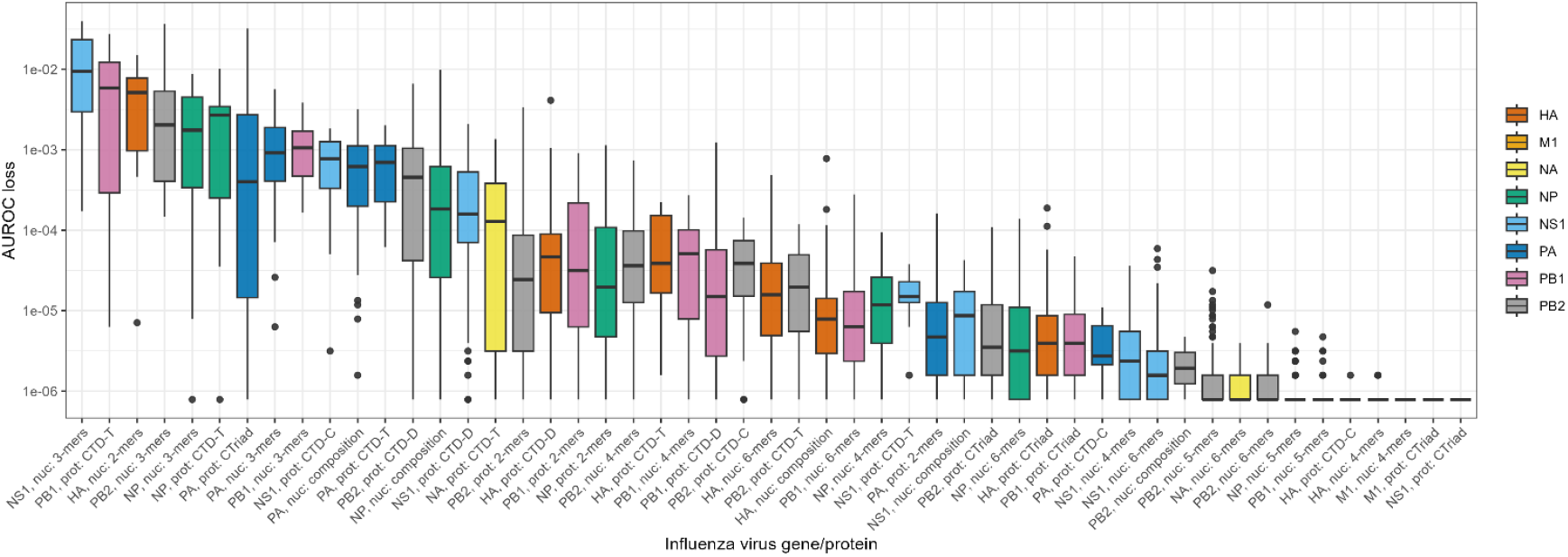
Permutation variable importance of avian influenza virus gene/protein features. Variable importance of feature sets capturing characteristics of avian influenza virus nucleotide or protein sequences, measured by reduction in performance (loss in AUROC) when individual features were permuted. Features are grouped together by feature set-gene or feature set-protein combinations into boxplots with outliers. Only feature sets that were retained in at least one stack ensemble are depicted. Boxes are ordered by median performance loss, shown on a log scale. Colour denotes gene/protein.

## Discussion

Here, we used a novel training strategy to build ensembles of machine learning models capable of predicting zoonotic capability purely based on avian influenza genome sequences. We find physiochemical features of proteins and also certain nucleotide motifs to be informative of zoonotic potential. While we report slightly lower performance than previous zoonotic prediction models for avian influenza (Eng et al. 2017, Scarafoni et al. 2019), we assert our stack model approach to be more generalisable to phylogenetically distant sequences as it achieves effective performance on entirely unseen influenza subtypes, including rare subtypes.

We achieved this by curating a representative training set informed by sequence similarity (Fig. 1) to ensure models would be robust against phylogenetic biases that pervade sequence-based machine learning (Mollentze and Streicker 2023). This also allowed us to intuitively handle a second challenge common to host predictor models: determining ground truth negatives for zoonotic infection. Since sequence metadata do not contain epidemiological context, host predictor models often resort to treating sequences as negative for zoonotic potential based purely on being sampled from a non-human host (Borkenhagen et al. 2021). Sequence clustering allowed us to conduct dissimilarity-based sampling to select an appropriate training set of negatives, ensuring model generalisability.

Following stack model training, optimal zoonotic predictions were made by using information from all but one segment. This suggests that signals of zoonotic potential are widespread throughout the influenza virus genome, beyond surface proteins and polymerase proteins – a finding consistent with previous influenza host predictor models (Eng et al. 2017). The exception was matrix protein M1, which had no functional influence on stack predictions (Fig. 4), despite individual models trained on this protein achieving good performance (Fig. 2). This suggests zoonotic signal is present in M1 but is redundant when considering features in other genes and proteins. Neuraminidase features also gave the lowest performance in individual models (Fig. 2) and consequently, only minimally contributed to final stack predictions (Fig. 4).

Collectively, non-zoonotic and zoonotic sequences were highly separable in predicted probability value (Fig. 3), with most zoonotic sequences being correctly predicted. The exceptions were among some zoonotic sequences of H7N9 which showed high intra-subtype variation in prediction, potentially reflecting the genetic diversity across separate “waves” of human infection (Quan et al. 2018), plus two sequences from isolated H7N3 spillovers (*A/Mexico/InDRE7218/2012* and *A/Canada/rv504/2004*). Among predictions for avian-origin sequences, we found a small number of H4 influenza viruses to have elevated zoonotic risk compared to other viruses of the same subtypes. Although no zoonotic spillovers of H4 avian influenza have been reported, there are suggestions these viruses may have zoonotic potential from serological evidence in exposed individuals (Kayali et al. 2010), albeit with potential for cross-reactivity.

### Relation to molecular mechanisms

While machine learning approaches have clear value in prediction, the fundamental importance of learning new biological insights from these methods is often overlooked (Borkenhagen et al. 2021). We report consistent zoonotic signal in influenza virus polymerase (PA, PB1, PB2), haemagglutinin (HA), nucleoprotein (NP), and non-structural protein (NS1) that could hint towards interaction mechanisms between viruses and host cells.

Mutations most closely associated with initial spillover to mammals tend to be those in polymerase genes that arise in avian populations or early in mammalian adaptation, particularly in the PB2 gene which includes the strongly mammalian-characteristic E627K mutation (Capelastegui and Goldhill 2025). This aligns with the generally greater importance of polymerase features than haemagglutinin features we observe (Fig. 4), as hemagglutinin traits act as a later adaptive barrier (Long et al. 2019). We find a mixture of k-mer, compositional, and protein features of PA, PB1, and PB2 to strongly influence zoonotic predictions (Fig. 4, Table 1). This could suggest changes to polymerase that enhance activity or compatibility with the host ANP32 protein family needed for viral replication, with the ANP32A homologue being distinctly shorter in mammals than birds (Capelastegui and Goldhill 2025).

For haemagglutinin, we find dinucleotides to have high variable importance in our model, with other feature representations having minor importance (Fig. 4). The primary trait of HA associated with host suitability is the efficiency of binding sialic acid, with α2,6-linked sialic acid being more prominent in human upper airway versus α2,3-linked sialic acid, which is more prominent throughout the avian gastrointestinal tract (Suttie et al. 2019). However, α2,6 preference is not a strict requirement for zoonotic infection, as α2,3-linked sialic acids also occur in human lower airways, potentially explaining the more modest influence of HA protein features within the stack.

Features of NP were also heavily relied on to distinguish zoonotic sequences. NP proteins of avian influenza viruses are targeted by multiple arms of innate immunity; human MxA proteins prevent nuclear entry of viral ribonucleoprotein complexes and human BTN3A3 proteins restrict transcription (Pinto et al. 2023). The features we identify could indicate NP protein motifs more likely to evade targeting and successfully establish human infection.

Most surprisingly, we report the most influential feature set captured NS1 nucleotide patterns (Fig. 4, Table 1). Although some fitness-increasing mutations in NS1 are known, likely via inhibition of host interferon responses, NS1 mutations have generally not been indicative of any specific host species adaptation (Long et al. 2019). As this feature set reflected k-mers across the entire segment, this may be capturing dynamics of mRNA splicing, which can produce the alternative protein NEP. The relative ratio of NEP:NS1 splicing products appears associated with human cell compatibility of avian influenza viruses (Long et al. 2019).

### Caveats and future directions

We note that sequence-based machine learning to predict viral traits carries some limitations. Supervised learning algorithms consider data to be both independent and representative and unlike phylodynamic approaches, cannot therefore infer chains of transmission or directional evolution. Although we introduced a pre-processing step to resample sequences and reduce the influence of phylogenetic bias upon models, we also cannot rule out some learned patterns being drawn from phylogeny via similarity in feature sets between related sequences. We also acknowledge that the genomic and proteomic features we use do not capture more complex molecular mechanisms like variation in post-translational modification such as glycosylation (Kim et al. 2018) that are likely to further affect cellular host compatibility.

More comprehensive sets of sequences for model training that capture wider variation in avian influenza and missing intermediates between avian and zoonotic sequences would benefit future models, though detecting zoonotic infections is problematic for asymptomatic cases. Alternatively, in-vitro cell line studies with a library of different influenza viruses or proteins in isolation (e.g., via lentiviral pseudotyping (Dufloo et al. 2025)) could generate wider training data coverage, at least for predicting human cell entry.

Among the model algorithms we used, random forests and XGBoost gave the most consistent performance. Both are tree-based methods which can represent highly structured, non-linear interactions between input features and are thus well-suited for large sets of genomic traits that individually have only very subtle effects on zoonotic potential. Newer computational advances to detect complex, subtle signals in unstructured data are rapidly developing in the field of deep learning. This includes protein language models (PLMs), large and densely interconnected neural networks capable of learning the ‘rules’ governing biological validity of nucleotide and protein sequences. Successful predicting zoonotic potential of influenza viruses using PLMs has so far proven elusive (Kawasaki et al. 2025), though promising new PLMs tailored exclusively to virus genomics are now emerging (Pan et al. 2025).

### Conclusions

The high accuracy of our stack model in predicting zoonotic potential of unseen influenza virus sequences suggests potential uses of these methods in practice. Predictions from our model (among others) can act as a rapid and inexpensive method to risk-assess newly emerging lineages of avian influenza for zoonotic spillover, requiring only a genome sequence as input. Careful interpretation of these model predictions in context could also be used for early detection of potential pre-adaptive mutations arising as influenza circulates through bird populations. For new lineages flagged as having high zoonotic potential, our methods could also triage choice of candidates for pre-emptive vaccine development, which could even include vaccines for poultry as an occupational health protection measure. We assert that machine learning and artificial intelligence systems can play an active role in pandemic prevention – but they must be trained carefully, putting the biology of host-virus interactions first and foremost.

## Acknowledgements

We would like to thank Gytis Dudas, Spyros Lytras, Rubaiyea Farrukee and Patrick Reading for providing expert comments that helped develop this work. We would also like to thank Lorenzo Cattarino, Rachel Abbey, Charlotte Belsey, and Juan Adriano within the Data Science & Geospatial Division of Health Analytics and Automation at the UK Health Security Agency for providing technical assistance and code validation.

This work made use of the Barkla High Performance Computing facilities at the University of Liverpool, UK. LB, JM, and MB would like to acknowledge funding support by CSL Seqirus UK Ltd to the University of Liverpool through the 2022 – 2027 Framework Research Collaboration Agreement Influenza Research Partnership, collaboration plan 002.

## Data and Code Availability

All code used to process data and generate analyses is available at https://github.com/lbrierley/ai_for_ai/

Machine learning feature datasets and final trained machine learning models are available at https://doi.org/10.5281/zenodo.17068424

