## Supplemental Table and Figures for "An AI for an AI: identifying zoonotic potential of avian influenza viruses via genomic machine learning"

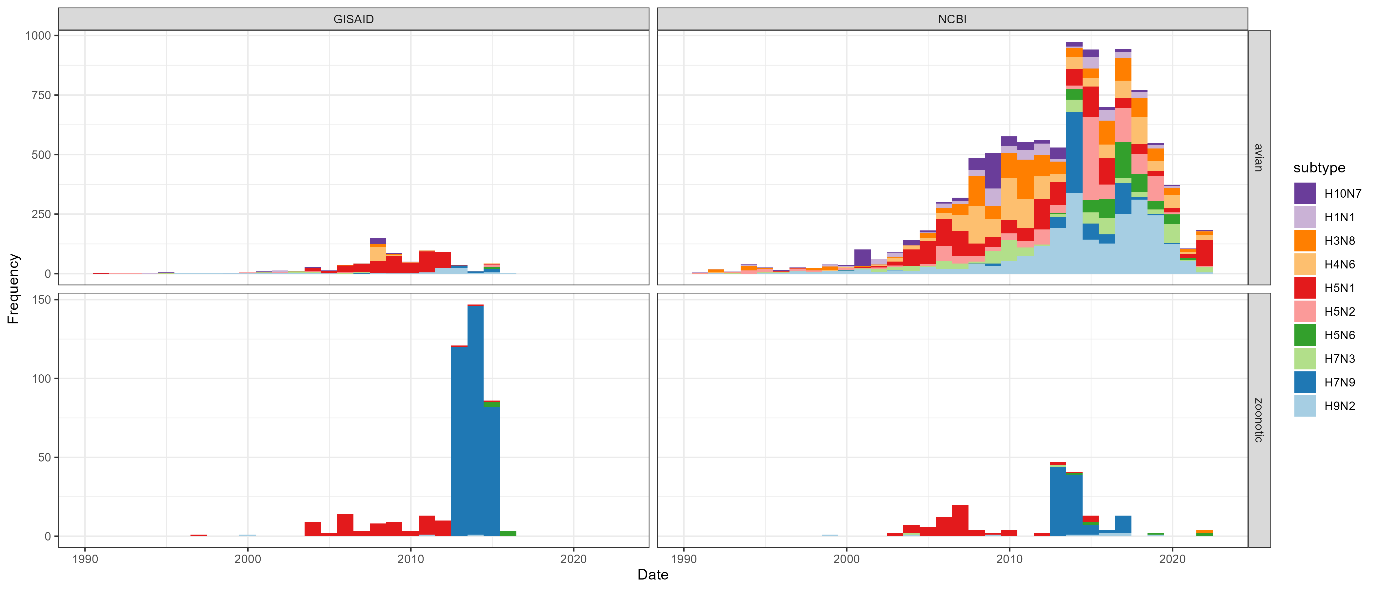
**Supplemental Figure S1. Temporal distribution of whole genome sequences of avian influenza virus.**
Stacked barplots of annual frequency of dated avian influenza virus whole genome sequences from 1990 onwards in filtered dataset before cluster downsampling. Barplots are separated by data source (GISAID and NCBI Influenza Virus Resource) and sample host (avian or zoonotic). Colours indicate subtype (only ten most common subtypes shown). Where sequences were deposited to both databases, only NCBI record was retained.

**Supplemental Table S1. Full subtype representation of whole genome sequences of avian influenza virus.**
Number of extracted avian influenza virus whole genome sequences in filtered dataset before cluster downsampling. Shading separates subtypes with at least one zoonotic sequence, and underline denotes subtypes selected a priori as holdout test sets. Table is sorted by frequency of non-zoonotic sequences for exclusively avian subtypes, and by frequency of zoonotic sequences for zoonotic subtypes.

| **Subtype** | **Non-zoonotic** | **Zoonotic** | **Subtype** | **Non-zoonotic** | **Zoonotic** | **Subtype** | **Non-zoonotic** | **Zoonotic** |
| --- | --- | --- | --- | --- | --- | --- | --- | --- |
| **H8N1** | 1 | 0 | **H4N5** | 13 | 0 | **H2N2** | 67 | 0 |
| **H9N4** | 1 | 0 | **H5N7** | 13 | 0 | **H1N3** | 73 | 0 |
| **H10N0** | 1 | 0 | **H2N4** | 15 | 0 | **H1N2** | 74 | 0 |
| **H15N4** | 1 | 0 | **H4N1** | 15 | 0 | **H6N5** | 76 | 0 |
| **H3N4** | 2 | 0 | **H2N8** | 16 | 0 | **H13N2** | 82 | 0 |
| **H8N8** | 2 | 0 | **H3N5** | 16 | 0 | **H10N5** | 98 | 0 |
| **H14N2** | 2 | 0 | **H4N4** | 16 | 0 | **H13N8** | 110 | 0 |
| **H14N7** | 2 | 0 | **H12N1** | 16 | 0 | **H7N1** | 122 | 0 |
| **H14N8** | 2 | 0 | **H12N2** | 16 | 0 | **H5N3** | 123 | 0 |
| **H8N2** | 3 | 0 | **H3N7** | 21 | 0 | **H10N3** | 124 | 0 |
| **H13N3** | 3 | 0 | **H4N3** | 21 | 0 | **H11N2** | 129 | 0 |
| **H5N4** | 4 | 0 | **H12N3** | 22 | 0 | **H7N2** | 151 | 0 |
| **H9N3** | 4 | 0 | **H2N5** | 26 | 0 | **H8N4** | 152 | 0 |
| **H9N6** | 4 | 0 | **H14N3** | 26 | 0 | **H4N2** | 169 | 0 |
| **H14N5** | 4 | 0 | **H3N3** | 27 | 0 | **H3N6** | 198 | 0 |
| **H2N6** | 5 | 0 | **H9N5** | 28 | 0 | **H13N6** | 207 | 0 |
| **H9N8** | 5 | 0 | **H5N9** | 29 | 0 | **H12N5** | 227 | 0 |
| **H14N6** | 5 | 0 | **H9N1** | 29 | 0 | **H6N8** | 237 | 0 |
| **H1N4** | 6 | 0 | **H1N6** | 30 | 0 | **H4N8** | 238 | 0 |
| **H3N0** | 6 | 0 | **H7N8** | 32 | 0 | **H2N3** | 246 | 0 |
| **H6N7** | 6 | 0 | **H11N8** | 33 | 0 | **H16N3** | 265 | 0 |
| **H12N6** | 6 | 0 | **H1N8** | 34 | 0 | **H5N8** | 338 | 0 |
| **H15N5** | 6 | 0 | **H9N9** | 34 | 0 | **H3N2** | 371 | 0 |
| **H12N9** | 7 | 0 | **H10N9** | 38 | 0 | **H6N1** | 397 | 0 |
| **H14N4** | 7 | 0 | **H4N9** | 40 | 0 | **H6N6** | 408 | 0 |
| **H15N9** | 7 | 0 | **H2N1** | 41 | 0 | **H11N9** | 425 | 0 |
| **H1N7** | 8 | 0 | **H12N4** | 41 | 0 | **H6N2** | 563 | 0 |
| **H4N7** | 8 | 0 | **H5N5** | 42 | 0 | **H1N1** | 569 | 0 |
| **H12N7** | 8 | 0 | **H2N7** | 46 | 0 | **H10N7** | 632 | 0 |
| **H6N4** | 9 | 0 | **H10N2** | 46 | 0 | **H5N2** | 1220 | 0 |
| **H6N9** | 9 | 0 | **H11N1** | 47 | 0 | **H4N6** | 1397 | 0 |
| **H7N5** | 9 | 0 | **H13N9** | 49 | 0 | **H7N4** | 25 | 1 |
| **H9N7** | 9 | 0 | **H10N6** | 50 | 0 | **H7N7** | 242 | 1 |
| **H11N7** | 9 | 0 | **H1N9** | 55 | 0 | **H3N8** | 1335 | 2 |
| **H11N6** | 11 | 0 | **H7N6** | 56 | 0 | **H7N3** | 702 | 2 |
| **H1N5** | 12 | 0 | **H2N9** | 57 | 0 | **H10N8** | 95 | 2 |
| **H6N3** | 12 | 0 | **H10N1** | 59 | 0 | **H5N6** | 499 | 12 |
| **H11N5** | 12 | 0 | **H3N1** | 61 | 0 | **H9N2** | 2372 | 13 |
| **H12N8** | 12 | 0 | **H11N3** | 63 | 0 | **H5N1** | 1832 | 137 |
| **H3N9** | 13 | 0 | **H10N4** | 65 | 0 | **H7N9** | 696 | 448 |

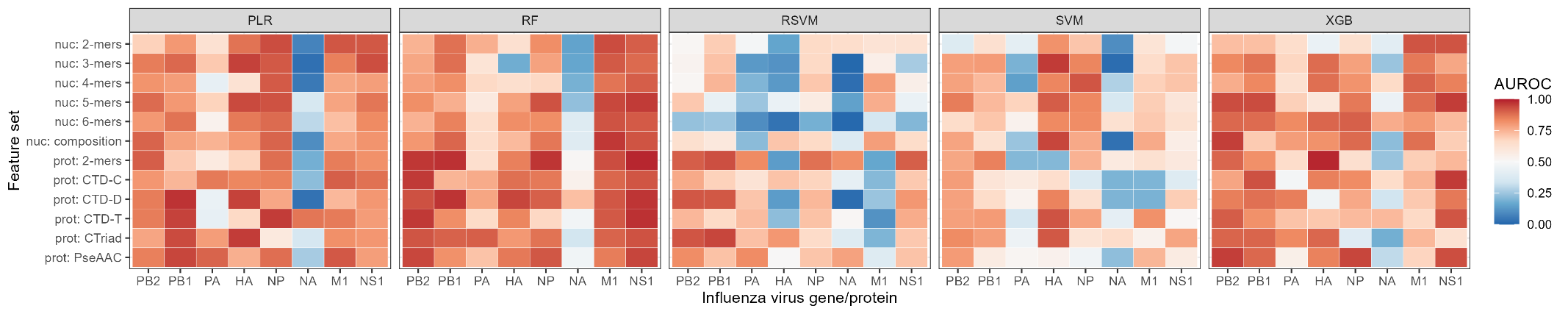
**Supplemental Figure S2. AUROC for individual segments, feature set, and algorithm combinations.**
Heatmap of performance for machine learning models covering each combination of influenza virus gene or protein (columns), sequence-derived feature sets (rows), and model algorithms (panels). Colour scale denotes collective performance over each of the thirteen held-out test subtypes aggregated into a single measure of AUROC. ‘nuc’ denotes nucleotide sequence features, ‘prot’ denotes protein sequence features: dipeptide composition (2-mers), Composition-Transition-Distribution (CTD), Conjoint Triad (CTriad), and Pseudo-Amino Acid Composition (PseAAC). PLR = penalised logistic regression, RF = random forest, RSVM = radial support vector machine, SVM = linear support vector machine, XGB = XGBoost.

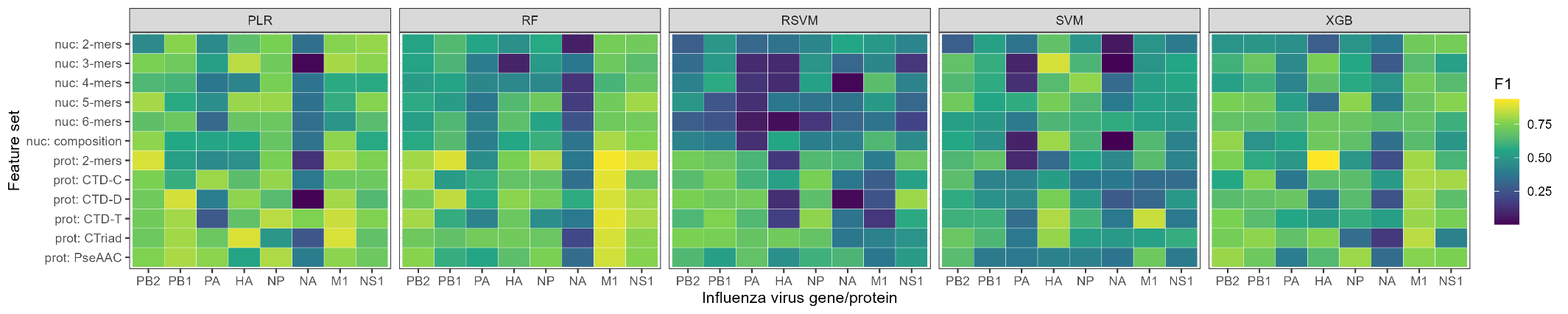

**Supplemental Figure S3. F1 score for individual segments, feature set, and algorithm combinations.**
Heatmap of performance for machine learning models covering each combination of influenza virus gene or protein (columns), sequence-derived feature sets (rows), and model algorithms (panels). Colour scale denotes collective performance over each of the thirteen held-out test subtypes aggregated into a single measure of F1 score. ‘nuc’ denotes nucleotide sequence features, ‘prot’ denotes protein sequence features: dipeptide composition (2-mers), Composition-Transition-Distribution (CTD), Conjoint Triad (CTriad), and Pseudo-Amino Acid Composition (PseAAC). PLR = penalised logistic regression, RF = random forest, RSVM = radial support vector machine, SVM = linear support vector machine, XGB = XGBoost.

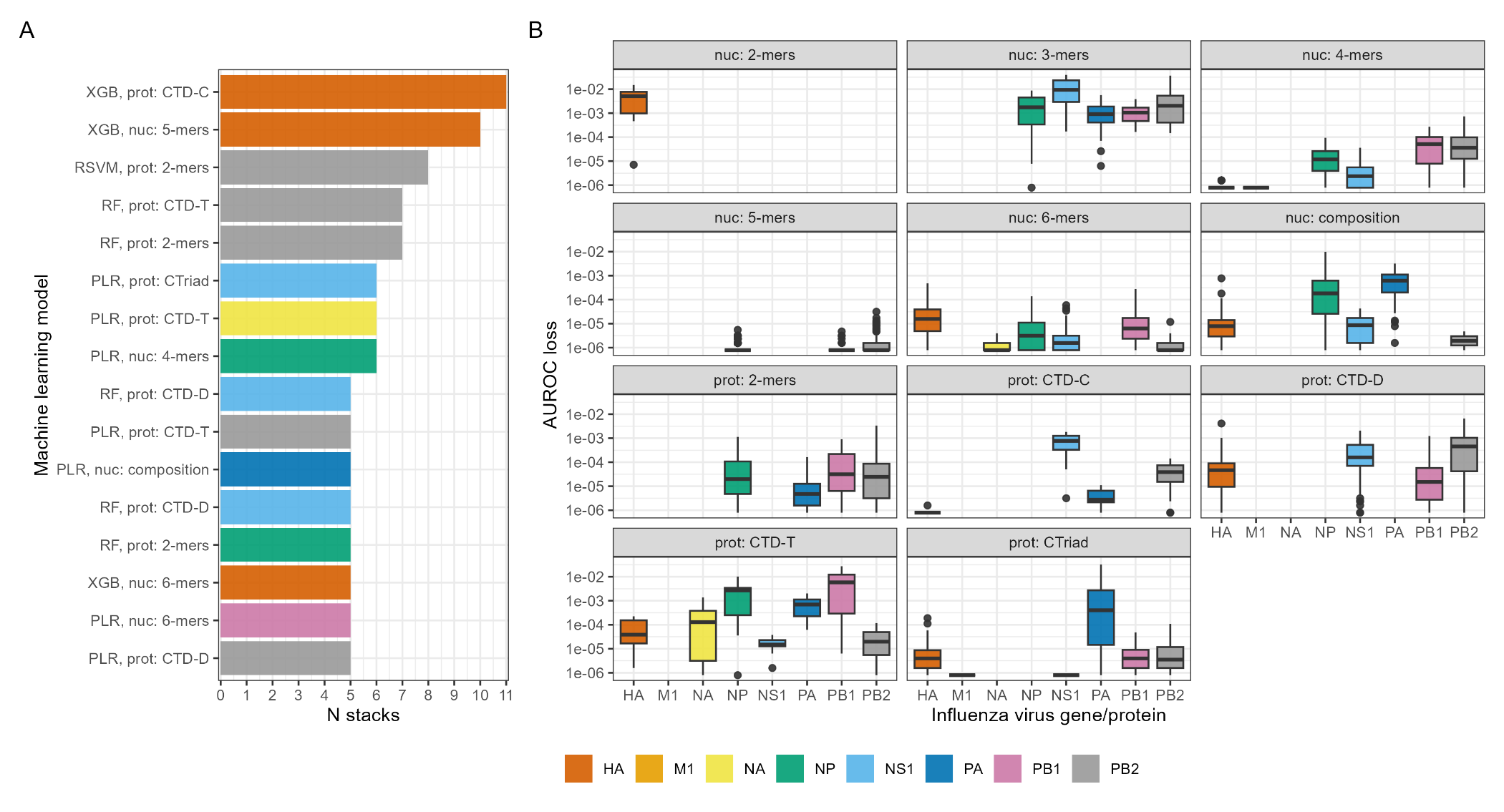
**Supplemental Figure S4. Importance of individual models in stack ensembles.**
Importance of feature sets capturing characteristics of avian influenza virus nucleotide or protein sequences. A) Retention of individual feature set-gene model combinations within stack ensembles, labelled with model method and feature set. Colour denotes gene/protein. Bars denote number of stack ensembles each model was retained in after model selection, of a possible maximum of thirteen (one stack ensemble is constructed omitting each of the held-out subtypes in turn), with only those in at least five stacks depicted. PLR = penalised logistic regression, RF = random forest, RSVM = radial support vector machine, XGB = XGBoost. B) Reduction in performance (log-scaled loss in Area under receiver operating curve) when individual features were permuted. Features are grouped together by feature set-gene or feature set-protein combinations into boxplots with outliers as in main Figure 4, with gaps representing models not retained in any stack ensembles after model selection.
